# Readiness Potential Prevalence During a Deliberate Decision-Making Task

**DOI:** 10.1101/2021.06.10.447898

**Authors:** J. Blignaut, D.J. van den Heever

**Affiliations:** Stellenbosch University

**Keywords:** decision-making, readiness potential, P300 potential, trauma, deliberate decision-making, electroencephalography

## Abstract

This study investigated the hypothesis that neural markers associated with arbitrary decision-making are present in higher order, deliberate decisions. Furthermore, the study aimed to investigate the effect of higher order decision content on neurophysiological markers such as the readiness potential and the P300 potential. An experiment was designed to measure, evaluate, and compare these electroencephalographic potentials under both arbitrary and deliberate choice conditions. Participants were presented with legal cases and had to convict and acquit criminal offenders. Distinct readiness potentials and P300 potentials were observed for both arbitrary and deliberate decisions across all participants. These findings support the hypothesis that the readiness potential and the P300 potential are present in the neurophysiological data for higher order deliberate decisions. The study also showed initial findings of how the readiness potential may inherently relate to decision content. Increased readiness potential amplitudes were observed for participants with previous exposure to violent crime when they had to acquit or convict criminals accused of violent crimes.

## 1. Introduction

Decision-making forms a fundamental part of executive cognition. Our lives are a series of choices: some are simple, while others require more deliberation. Unravelling the cortical networks that underlie the decision-making process plays an integral part in understanding consciousness and volition. Some past studies suggest that our choices are deterministically formed up to several seconds prior to conscious awareness of intent (Libet, et al., 1983; Soon, et al., 2008; Soon, et al., 2013). For decision-making studies, a distinction is typically made between voluntary and involuntary actions. The current study aimed to investigate the neural correlates underlying voluntary decision-making by measuring and investigating the neurophysiological differences and similarities between arbitrary and deliberate decisions. The aim was to show that the readiness potential (RP), as well as the P300 potential, is present in both cases. Moreover, the study aimed to investigate to what extent the content of higher order decisions affects the neurophysiological makers.

In 1983, Libet et al. designed an electroencephalographic (EEG) experiment wherein five participants were asked to act on the urge to flex the wrist of their dominant hand while reporting on the moment of awareness of intent (Libet, et al., 1983). Each participant completed six sessions of 40 trials for which the data were averaged to produce a trend approximating Kornhuber and Deecke’s RP. Even though no reported “preplanning” occurred, the recorded EEG-data showed a clear spike in neural activity 350 ms before the reported urge to move (i.e. moment of conscious choice), and 550 ms prior to movement (i.e. moment of action). Libet et al. used this finding as basis to question the notion of conscious will. They reasoned that the rise of the RP – observable 350 ms prior to awareness of intent in this “free” self-initiated task – supported their argument that free will is an illusory construct absent in self-initiated human action. Due to the contentious nature of the results, the findings of their study received extensive criticism. Critics argued that to act on the urge to flex a muscle cannot be considered a true measure of free choice (Wolpe & Rowe, 2014).

In 2008, Soon et al. conducted a similar experiment using functional Magnetic Resonance Imaging (fMRI). The experiment was adapted to include a choice task, thereby addressing one of the main criticisms of the original Libet study. The recorded neural responses enabled Soon et al. to predict the outcome of the choice, with relative accuracy, up to seven seconds prior to the participants’ reported subjective awareness (Soon, et al., 2008). Moreover, Soon et al. considered brain areas beyond the supplementary motor area (SMA) and pre-SMA to inform a more holistic understanding of the cortical networks underlying decision-making. In a later study, Soon et al. adapted their original experiment by increasing the complexity of the choice task, asking participants to add or subtract two numbers per choice trial (Soon, et al., 2013). Since then, the Libet and Soon experiments have been recreated for other EEG and fMRI studies with findings that support the original findings (Lavazza, 2016; Verbaarschot, et al., 2015). A different study by Alexander et al. found that the RP is present even in the absence of movement and that motor-related neural processes do not significantly affect the RP (Alexander, et al., 2016). Another study by Jo et al. corroborated these findings by setting up an experiment with a self-initiated movement condition as well as a no-movement condition (Jo, et al., 2013). They found that there was no significant difference between the movement condition RP and the no-movement condition RP. Herrmann et al. observed a clear RP build-up prior to stimulus presentation in a task where participants had to press one of two buttons depending on the stimulus presented (Herrmann, et al., 2008). Yet these existing studies were all limited to choosing between arbitrary alternatives.

The neurophysiological architecture that underlies deliberate decision-making has been mostly studied in the field of Neuroeconomics (Bossearts & Murawski, 2015). In 2019, Maoz et al. were first to introduce the concept of deliberate decisions into the neuroscience of human volition. They argued that the arbitrary decisions presented in previous studies were void of purpose, reason and consequence and that it therefore remains unknown to what extent the previous findings are applicable to decisions that matter (Maoz, et al., 2019). Maoz et al. defined deliberate decisions as decisions of interest, with ecological and real-life relevance. They developed an EEG choice task in which participants were instructed to donate money to one of two non-profit organisations (NPOs) (Maoz, et al., 2019). The experiment consisted of deliberate and arbitrary trials: for deliberate trials, the chosen NPO would receive a donation of $1000 and the NPO not chosen would receive $0. For arbitrary trials, regardless of the choice, both NPOs would receive an equal amount of $500. For arbitrary choice trials, clear RPs were observed while the deliberate choice trials were marked by an absence of RPs.

The current study further investigated the differences and similarities between arbitrary and deliberate decisions, with a specific focus on how subjective traumatic experiences may influence the neurophysiological markers of deliberate decision-making. For the purposes of this study, deliberate decisions were defined as choices with consequences.

## 2. Materials & Methods

### 2.1 Participants

Twenty-nine healthy participants (7 females; 22 males) aged 21 to 28 years, volunteered for the study. Three participants’ data were excluded from the study due to excessive noise and two participants were removed from consideration for being left-handed. Participants were recruited via email and institutional permission from Stellenbosch University was obtained to support the recruitment process. The experiment was approved by Stellenbosch University’s Health Research Ethics Committee (HREC) and was conducted in accordance with the ethical guidelines and principles of the international Declaration of Helsinki, South African Guidelines for Good Clinical Practice and the Medical Research Council (MRC) Ethical Guidelines for Research. Informed consent was obtained from all participants before taking part in the study. The sample of participants was further divided into different groups for cross-comparisons within the study.

### 2.2 Procedure

The study consisted of two parts, completed on two separate days. During the first part, participants were required to fill out a questionnaire illustrating their relationship to crime and violent crime throughout their lives. Since the study addressed issues with psychological valence, such as rape and murder, it was important to consider the respective participants’ histories with violent crime. Furthermore, since the study was conducted in South Africa, it was statistically likely that participants might have been affected by violent crime throughout the course of their lives – either directly or indirectly.

During the second part of the study, EEG data were collected while participants completed a choice task. Participants completed their EEG sessions individually. Upon arrival at the lab, participants were informed of the session’s proceedings and encouraged to ask questions if anything was unclear to them. Participants were also reminded that they could withdraw from the study at any point. For the EEG recordings, participants were asked to keep as still as possible and not to move their heads, or blink, excessively during the experiment. EEG noise due to eye movements while reading, were removed during data pre-processing (see Section 4.3). Participants were told to make themselves comfortable but to move only their corresponding index fingers as far as it was possible. Thereafter, the testing procedure was explained. Once the instructions had been communicated, participants were asked to assume their testing positions in front of the computer screen. All the verbal instructions were again presented in a written form. Following the instruction screens, participants were presented with a practice round during which they responded to three choice tasks. Once they understood how the practice round worked, testing began. Since the EEG setup made it possible to unplug the EEG cap, participants were also encouraged to take breaks. At the end of the lab session, everyone was remunerated for their participation.

The choice task asked participants to compare two different crimes, simultaneously presented on the left (criminal F) and the right (criminal J) sides of computer screen. They then had to choose who to convict/acquit for each scenario pair, i.e. per choice trial. All participants completed 360 choice trials, divided into 6 blocks of 60 trials each. Participants were randomly divided into two equal participant groups: one group had to decide whether to acquit criminal F or criminal J; the other group had to decide whether to convict criminal F or criminal J. They were told to place their index fingers of their left and right hands on the ridged “F” and “J” keys, respectively. The crimes presented contained the summarised case details of the competing offences (see Figure 1). The summaries provided enough detail for participants to make an informed decision, while still easy to read and understand. The crimes were reconstructed using the details of existing criminal cases. Participants had access to a sheet summarising relevant legal terminology, however, the cases presented were self-explanatory. The cases related to criminal scenarios such as theft, arson, murder, rape, assault, attempted crimes, and crimes committed in self-defence. For each choice trial, participants had 10 seconds to respond after which a different screen prompted them to make a choice. They were told to take their time while responding but they were also informed that the computer would prompt them to respond if they were unresponsive for too long. The prompt was to ensure that participants remained focused throughout the task. Figure 1 graphically illustrates the typical sequence of a single choice trial, along with the display times per frame.

**Figure 1:**
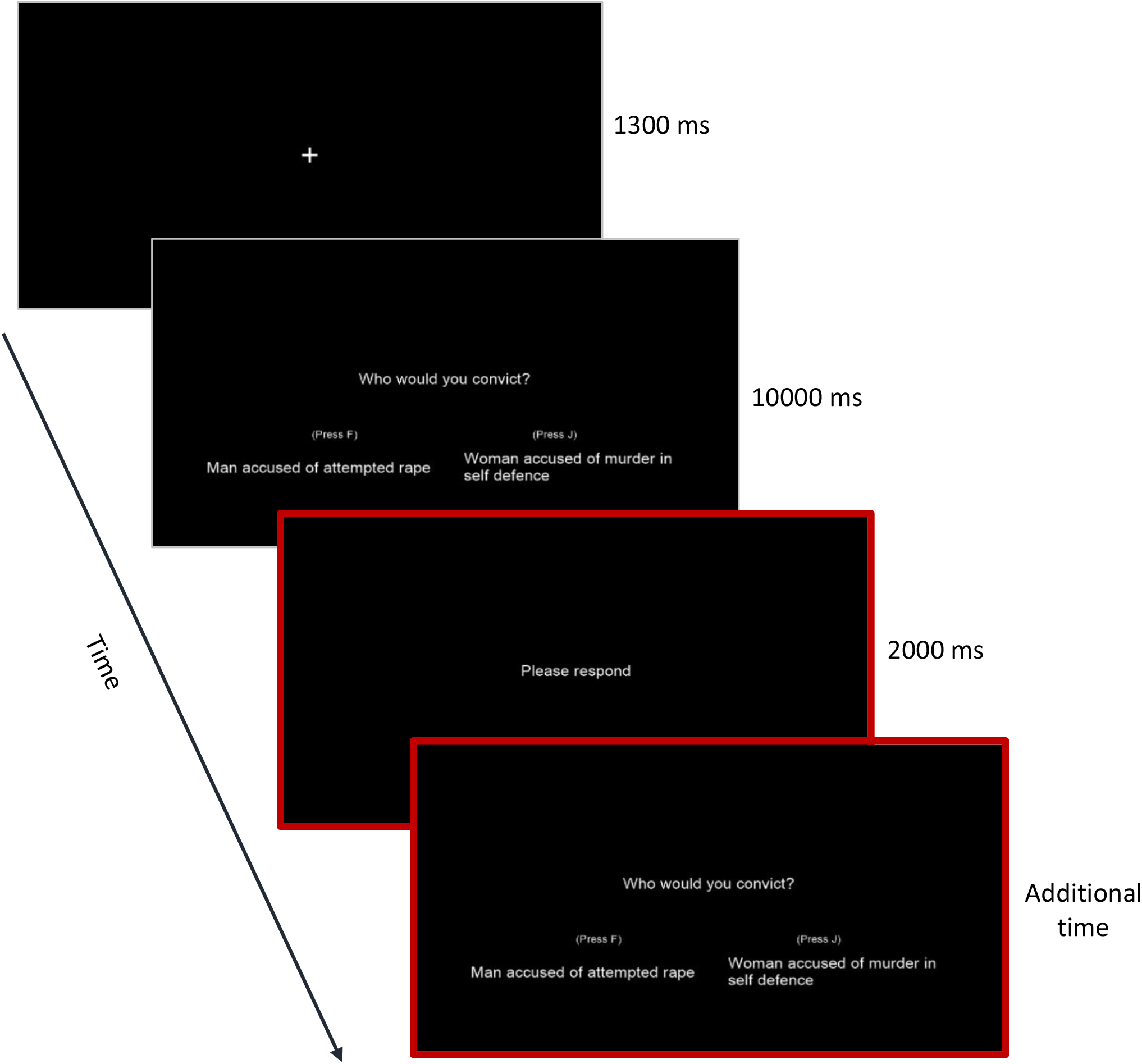
Typical sequence of a decision trial (the red frames indicate screens displayed if participants take longer than 10 s to make a choice)

In similar recent studies conducted by Maoz et al. and Verbaarschot et al., the researchers made a clear distinction between arbitrary and deliberate choices (Maoz, et al., 2019; Verbaarschot et al., 2019). This distinction served to separate the arbitrary choices found in previous Libet-type RP studies from deliberate decisions with real-world consequences. Consequently, the choice trials in this study were divided into two blocks: arbitrary and deliberate. At the start of each block, the programme informed participants whether they were responding to an arbitrary or a deliberate decision block. To eliminate potential experimental biases, participants performed the task blindly. They were told that the experiment was designed to evaluate whether EEG can be used to improve the jury selection process for prospective legal trials. This misdirection was necessary to maintain the integrity of the study, since knowledge of the deliberate decision-making investigative component may have influenced participants’ responses and biased the results (Bode, et al., 2014). Not knowing the full scope of the study did not pose any harm to participants. They believed the task was an audition for compiling a jury of 12 jurors for a mock legal trial and that their neurophysiological data collected during the EEG task would serve as selection criteria for the jury selection process. Furthermore, they were informed that for deliberate blocks, their responses would be evaluated for the jury selection process and that they were responding to unsolved cases; and for arbitrary blocks their responses would not be evaluated and they were responding to solved cases. These distinctions aimed to ensure that participants would consider evaluated/unsolved cases with more deliberation than non-evaluated/solved cases.

For the EEG recordings, a 128-channel Brain Products active channel amplifier (actiCHamp) EEG system (Brain Products, Germany) was used. For this study, 64 electrode channels were utilised - with a separate ground electrode channel located at FPz. The EEG data were sampled at 500 Hz, with a common reference located at Cz in the standard 10/20 system electrode positioning locations. The stimuli were presented on a 21-inch Dell monitor with a refresh rate of 60 Hz and a resolution of 1024×768 pixels. The experimental script was written using PsychoPy2 v1.90.2 (Peirce, 2009). Participants completed the experiment while sitting in a dimly lit, quiet room. All distractions, such as cell phones and smart watches, were removed for the duration of the experiment. All participants had normal, or corrected to normal, vision and were positioned 60 cm from the screen. The study was conducted at the Central Analytics Facility (CAF Unit), located at Stellenbosch University.

### 2.3 Data analysis

For the pre-processing of event-related potential (ERP) and EEG data, EEGLAB (a Mathworks MATLAB R2018a graphical user interface) was used. The recorded data were re-referenced to electrodes symmetrically placed across the scalp and located just inward of the mastoid electrodes (P7 and P8). Since Cz is a general area of interest when considering the RP, the mastoids are typically used as reference. For this study, the best results were obtained using linked mastoid (LM) referencing. However, since the data that was recorded from the typical mastoid sites, TP9 and TP10, was on average too noisy to produce reliable results, electrode sites P7 and P8 were used instead.

A digital finite impulse response (FIR) filter was used to filter the data between 1 and 40 Hz, using a band-pass filter. EEGLAB’s built-in independent component analysis (ICA) function, in conjunction with the multiple artefact rejection algorithm (MARA) plug-in, was used to remove all marked artefacts prior to analysis. Channels that were marked for removal, were replaced using interpolation. Data epochs were extracted from 3000 ms before to 350 ms after the button press event, and the data were time-locked to the button presses. Lastly, the epoched data were run through an artefact detection algorithm that eliminated trials with peak-to-peak amplitude differences exceeding 100 μV. This also ensured that noise due to excessive eye movements was removed.

The statistical analysis phase was divided into four parts. Firstly, a power calculation was performed on the data to determine the power of the study for a mean difference of 1.5 and a standard deviation of 1, with α = 0.05 and β = 0.2 (Noordzij, et al., 2010). The results showed that the sample size required for statistically significant difference with a power of 0.85, was roughly 10 participants (Table 1 shows the sizes of the different participant groups and subgroups compared in the study). Thereafter, descriptive univariate statistics was used to identify the mean, median, standard deviation, interquartile range, minimum and maximum values per variable. During univariate analyses, tests for normality were performed to determine which bivariate analysis procedure subsequently needed to be followed. Lastly, multivariate analyses determined the statistical significance between different participant subgroups. All statistical analyses were done using SAS (SAS Institute Inc., 2017), NCSS (Dawson & Trapp, 2004) and SAS Enterprise Miner (SAS Institute Inc., 2011).

**Table 1:** Different statistical power values for different participant groups (where the sample totalled 24 participants)

| Group | Sample size | Statistical power |
| --- | --- | --- |
| Acquit | 12 | 0.95 |
| Convict | 12 | 0.95 |
| Left button presses | 24 | 0.99 |
| Right button presses | 24 | 0.99 |
| Arbitrary | 24 | 0.99 |
| Deliberate | 24 | 0.99 |
| Crime I: Y | 6 | 0.65 |
| Crime I: N | 18 | 0.98 |
| Crime II: Y | 13 | 0.96 |
| Crime II: N | 11 | 0.92 |

## 3. Results

The data were divided into several different groups for comparison. Firstly, the differences between the acquit and convict trials were evaluated, then the differences between the left and right button press responses, and thereafter the differences between the deliberate and arbitrary blocks (see Table 1). Lastly, the information gathered from the participant questionnaires was considered. The questionnaires were designed to determine the respective participants’ relationship to violent crime. Based on these findings, participants were divided into separate groups where participants had personally been exposed to violent crime (Crime I); and where participants had close relatives who had been exposed to violent crime (Crime II). Each of the Crime I and Crime II groups contained Yes (Y) and No (N) subcategories, i.e. someone who, for example, had personally been exposed to violent crime would be in the “Yes” subcategory of the Crime I group (Crime I: Y) and someone who had not would be in the “No” subcategory of the Crime I group (Crime I: N). For the purposes of this study, violent crimes were defined as assault or sexual assault.

The parameters evaluated within the study were EEG scalp potentials, button press responses and response times. The relevant ERPs considered for analysis were the RP and P300 peaks. The statistical significance of the RP and P300 scalp potentials was evaluated using the 95% confidence interval (CI), while the statistical significance of the button press responses and participant response times were evaluated using the 95% CI, analysis of variance (ANOVA) and Wilcoxon ranked sum (WRS) tests. The button press event was taken as time zero (Maoz, et al., 2019). The P300 peak was taken as the maximum peak occurring at any position 250 to 350 ms post button press.

### 3.1 Acquit vs convict

No statistically significant difference was found comparing the response times for the acquit and convict participant groups (ANOVA = 0.03, p = 0.8587), i.e. there are no statistically significant differences between the red and yellow error bars of the two different participant groups shown in Figure 2. This finding demonstrates that participants considered acquit and convict trials without distinction. Additionally, there was no overlap in the 95% CI between the RP and P300 amplitudes of the acquit and convict trial types, indicating no statistically significant neurophysiological difference between the two trial types (see Figure 3). Subsequently, the data of the two participants groups were grouped together for further analysis.

**Figure 2:**
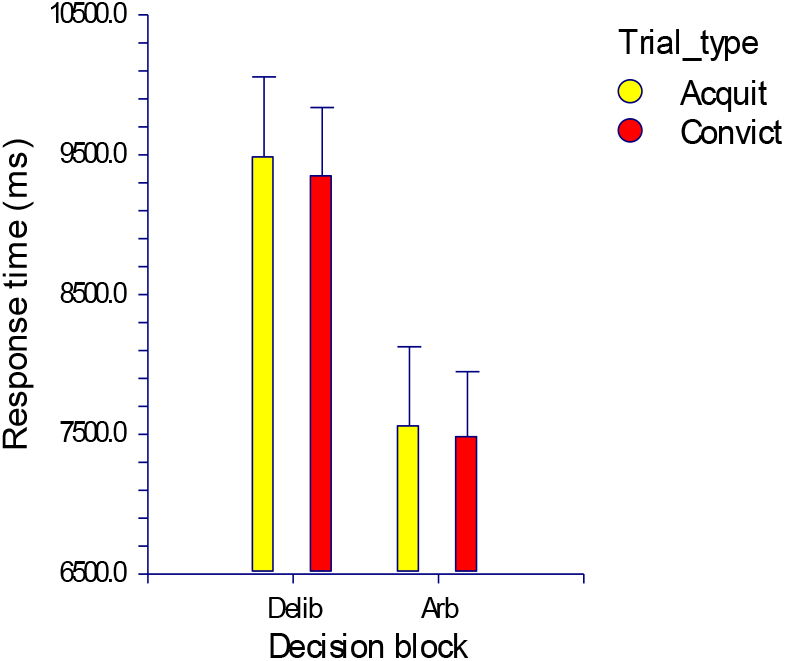
Error bars of response times for acquit and convict trials for deliberate (delib) and arbitrary (arb) decision blocks.

**Figure 3:**
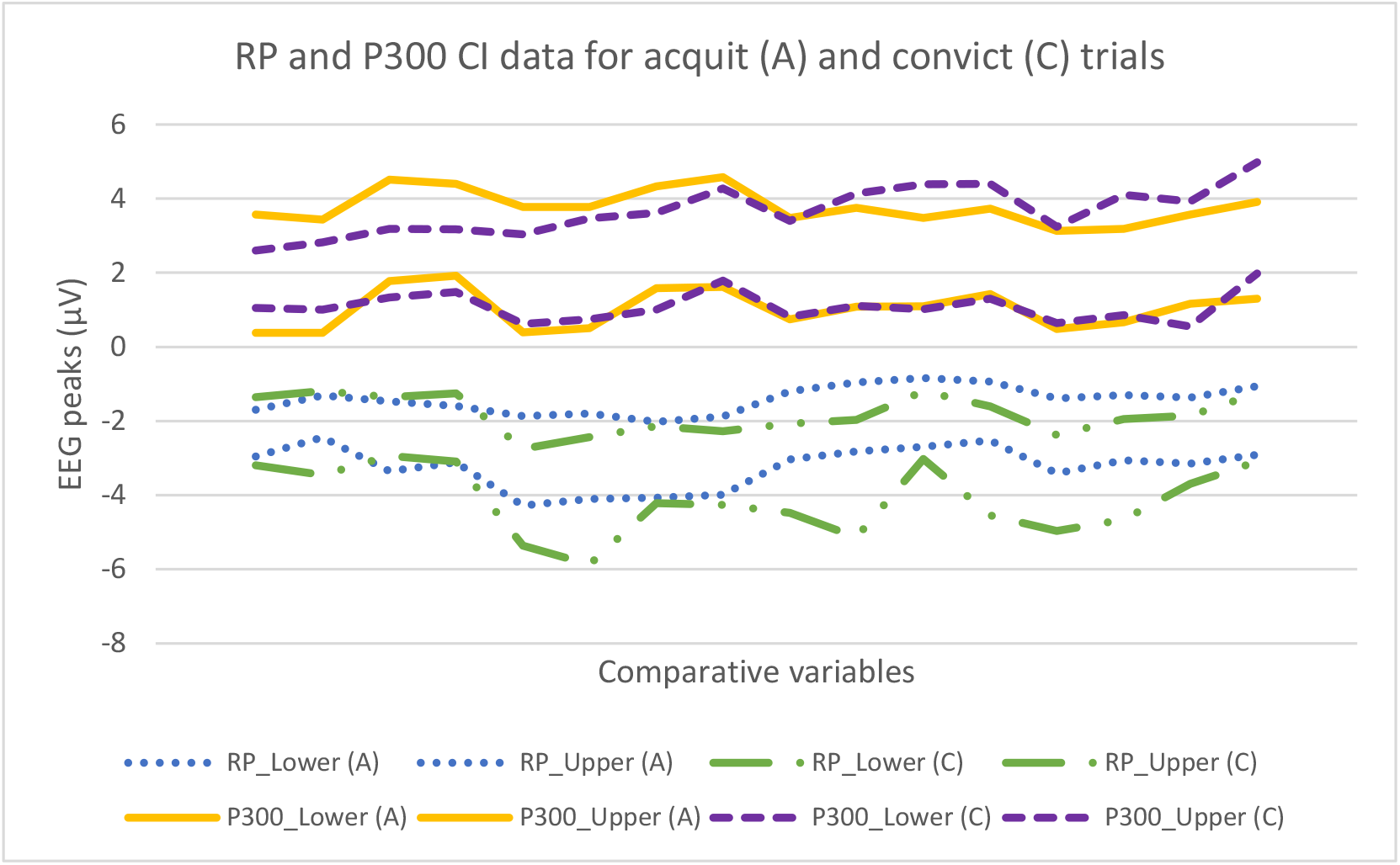
95% CI of the mean RP and P300 peaks for acquit and convict trials.

### 3.2 Left vs right

Similarly, comparing all participants’ left and right button press responses, the 95% CIs showed no statistically significant neurophysiological differences between left and right (see Table 2). Practically, this confirms that participants did not favour one hand over the other and the data was not skewed in favour of either hand’s button press responses. Subsequently, the button press responses were grouped to produce one set of results for further analysis.

**Table 2:** 95% CI of the mean RP and P300 peaks for left and right button press responses.

| Variable | Left CI (μV) |  | Right CI (μV) |  |
| --- | --- | --- | --- | --- |
|  | Lower 95% | Upper 95% | Lower 95% | Upper 95% |
| RP_Cz_arb | -2.81 | -1.79 | -2.69 | -1.51 |
| RP_Cz_delib | -2.84 | -1.71 | -2.81 | -1.71 |
| P300_Cz_arb | 1.08 | 2.72 | 1.09 | 2.73 |
| P300_Cz_delib | 1.92 | 3.48 | 2.03 | 3.45 |

### 3.3 Arbitrary vs deliberate

To compare the responses recorded during the arbitrary and deliberate decision blocks, the average response times and button press responses were considered. These parameters were used to statistically validate the initial distinction made between arbitrary and deliberate decisions. For this validation, the logic of the responses was plotted.

The trials were structured in a way that the questions in the convict trials exactly matched the questions in the acquit trials, for the two respective participant groups. Despite there being no wrong or right answers, in most cases the questions favoured one answer over another. Therefore, since the acquit and convict groups consisted of 12 participants each, it was expected that across all trials the average number of left button presses for the acquit trials should roughly match the average number of right button presses for the convict trials (see Figure 4).

**Figure 4:**
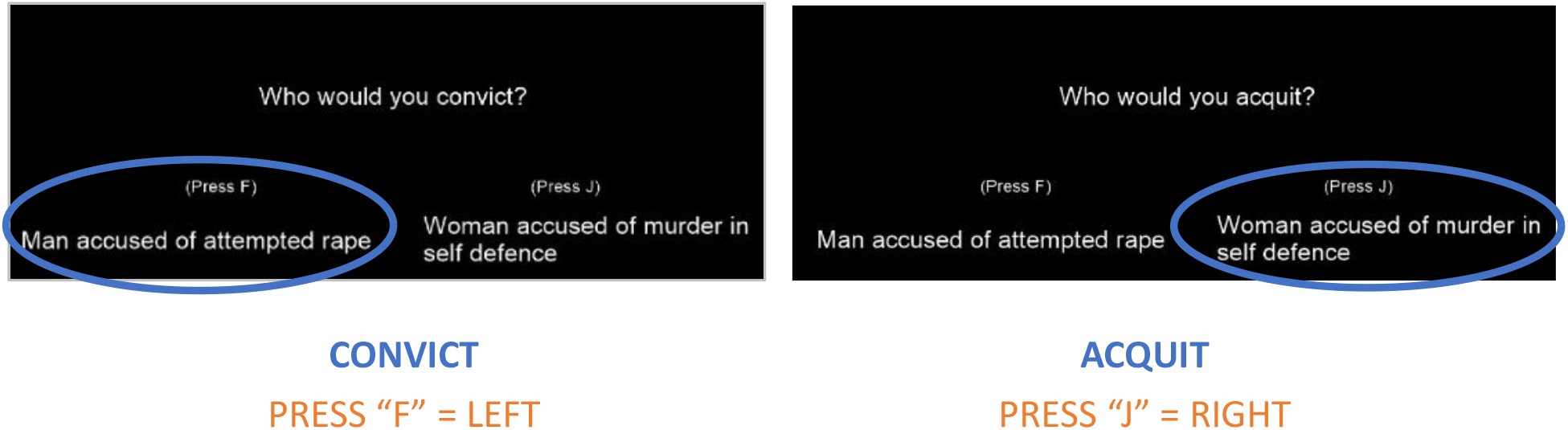
Comparative acquit and convict trials illustrating the expected logical outcome of the two trial types.

It was interesting to note that for the deliberate decision blocks this assumption proved correct. However, in the case of arbitrary decision blocks, there was a far less pronounced trend between the left button presses of one trial type and the right button presses of the other. Figure 5 graphically shows the cross-group similarity for deliberate blocks and discrepancy for arbitrary blocks. Looking at Figure 2, the response times also show a distinct difference between arbitrary and deliberate blocks. The 95% CIs of the response times confirm that participants responded significantly faster to arbitrary blocks than to deliberate blocks.

**Figure 5:**
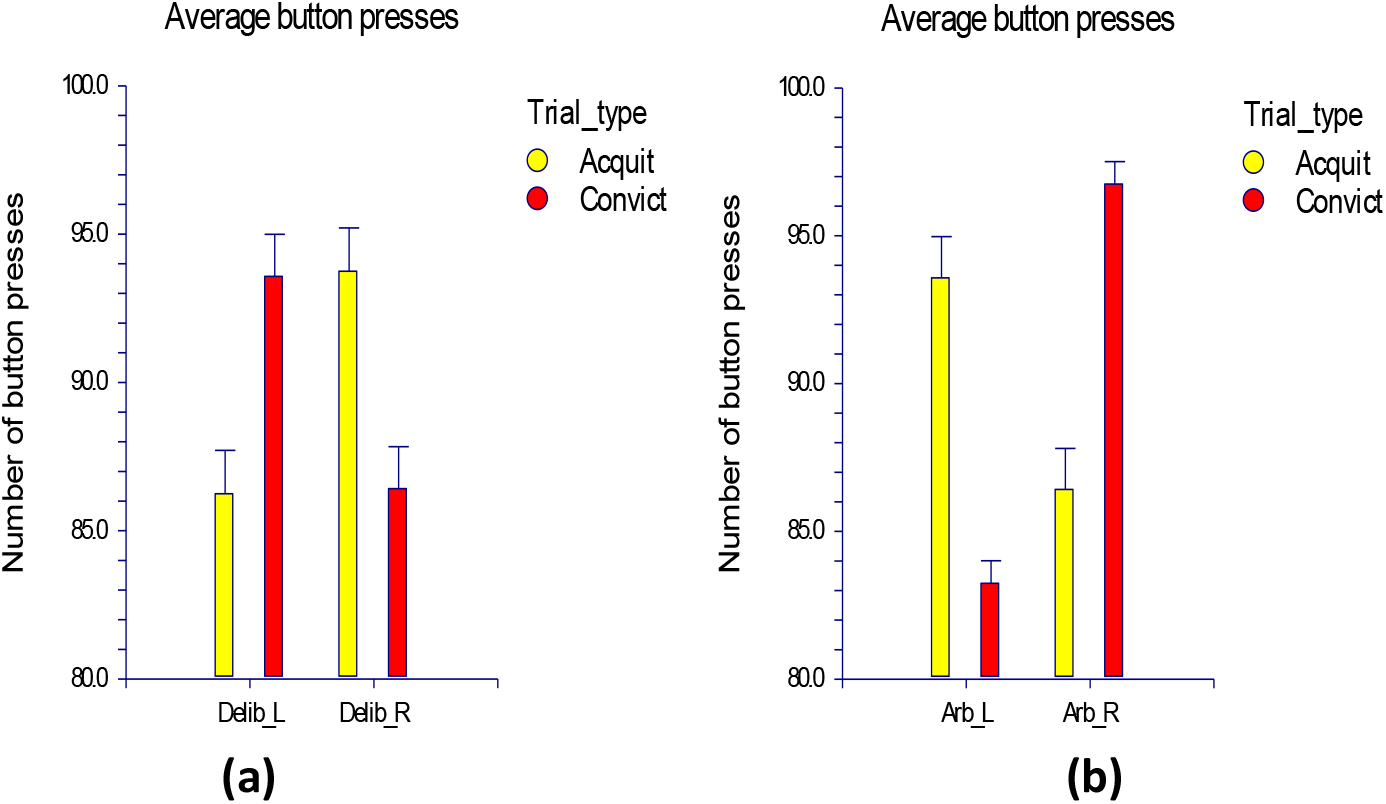
Number of button presses for a) deliberate (delib) and b) arbitrary (arb) decision blocks, for acquit and convict groups for all decision trials, averaged across all participants.

To evaluate and compare the ERPs for the different decision blocks, scalp data at electrode Cz were considered. Figure 6 shows a clear RP build-up for the arbitrary decision blocks, with an onset roughly 300 ms before the button press event. Figure 6 also shows a pronounced P300 peak 250 ms post button press. From Figure 6, the same RP build-up and P300 peaks are present for the deliberate decision blocks. This trend can be found for all participants across all trial types. Although the P300 peak was slightly attenuated in the case of arbitrary blocks, the RP trends between the two blocks were very similar. The black line in Figure 6 plots the difference between the two decision blocks. Looking at this line, the difference is minimal at the points of interest, i.e. where the RP and P300 peaks occur. The shaded grey areas in Figure 6 show regions of significance where the p-values were less than 0.05. The calculated difference fluctuates around zero, with unremarkably small null line deviations. Figure 6 shows the RP and P300 trends at electrode Cz. The same trends were found at electrode Fz, Fp1 and Fp2.

**Figure 6:**
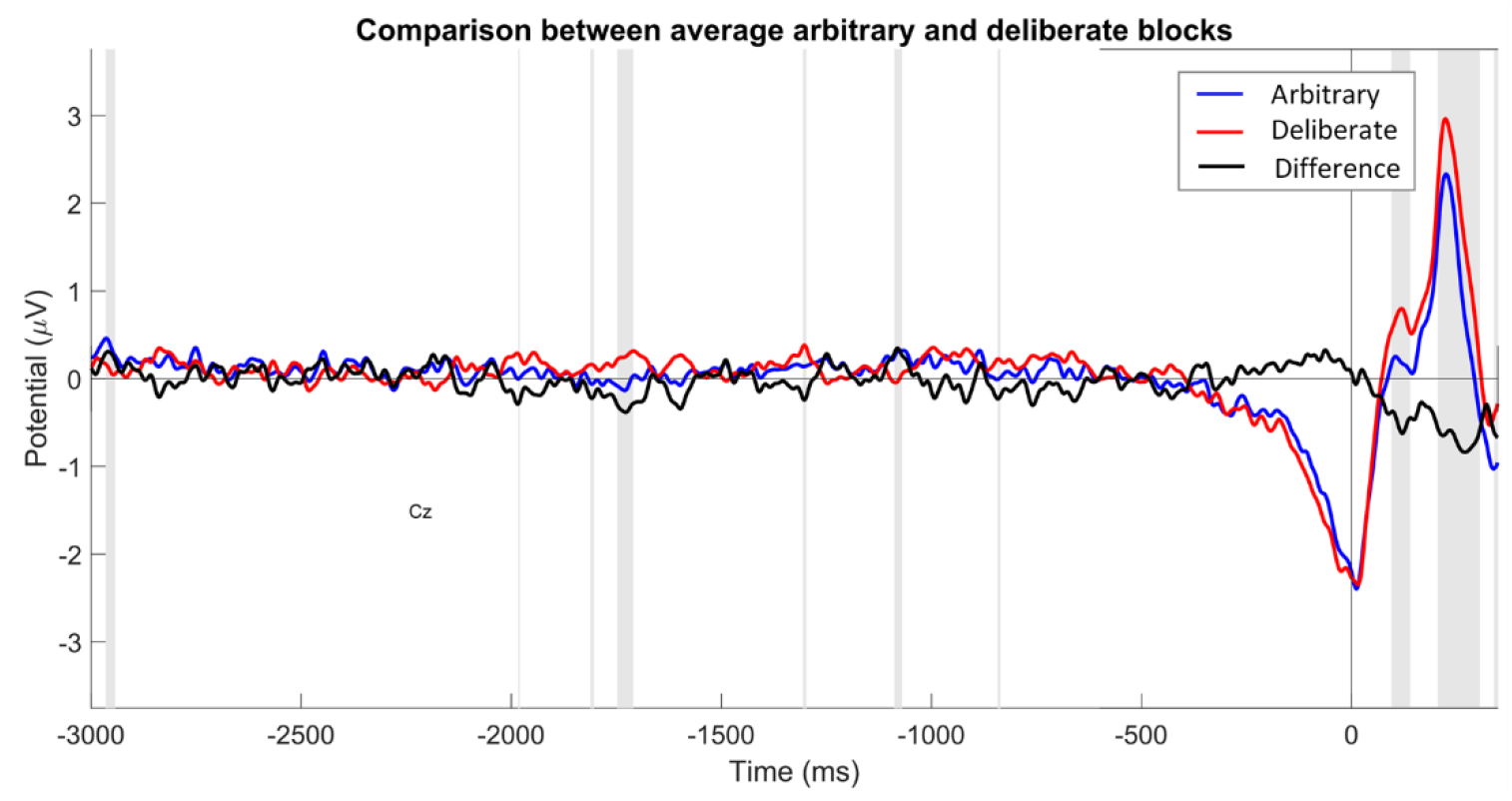
EEG scalp data comparison between RP and P300 peaks for arbitrary and deliberate decision blocks recorded from electrode position Cz.

Figure 7 illustrates the RP and P300 amplitudes for the Crime I group, i.e. participants with 1^st^ person exposure to violent crime (Crime I: Y) and participants with no 1^st^ person exposure to violent crime (Crime I: N). Since electrode Fz is responsible for recording activity from the intentional and motivational centres of the brain (Teplan, 2002), the Crime I comparative data is shown at electrode Fz. Looking at Figure 7, there are evident observable differences between the two groups, however the p-value differences were not statistically significant. The Crime I: Y and Crime I: N groups consisted of 6 participants (power = 0.65) and 18 participants (power = 0.98), respectively. The limited sample size for the Crime I: Y group may have influenced the statistical significance of the cross-group comparison. Yet, despite the lack of statistical significance, these results will still be discussed due to their potential relevance. Within the Crime I group, the RP amplitudes showed attenuation in the case of Crime I: N. Conversely, the P300 amplitudes showed attenuation in the case of Crime I: Y. The more pronounced RP amplitudes for Crime I: Y participants could suggest that the RP may be interrelated to decision content.

**Figure 7:**
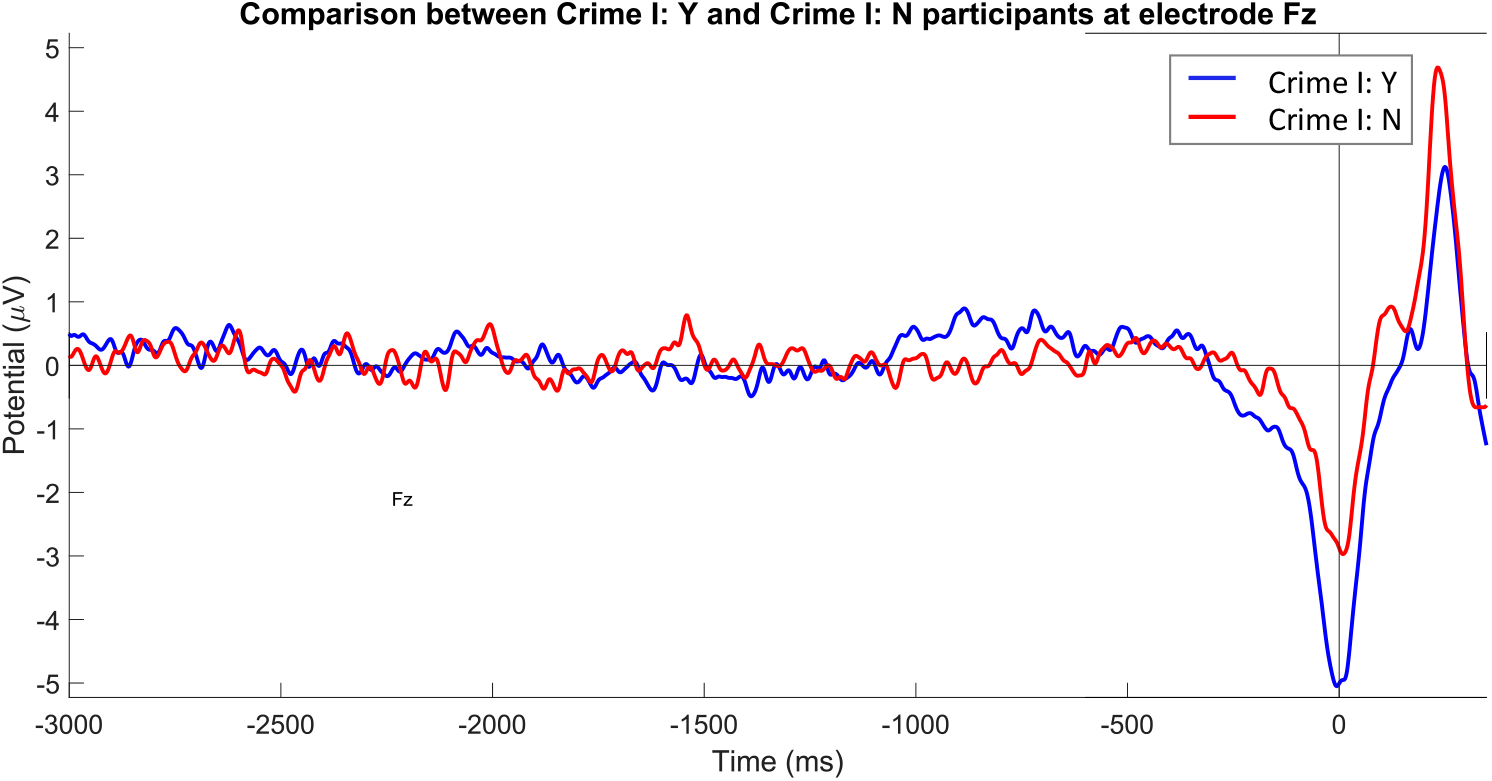
EEG scalp data comparison between 1^st^ person (Crime I: Y) and non-1^st^ person (Crime I: N) exposure to violent crime recorded from electrode position Fz.

## 4. Discussion

The aim of this study was to investigate the neurophysiological correlates underlying deliberate decision-making. This study defined two distinct decision classes, namely arbitrary and deliberate decisions. The experimental protocol aimed to investigate the neurophysiological differences between these two decision classes. Figure 2, Figure 4 and Figure 5 validate that participants considered deliberate and arbitrary blocks differently. The findings for the deliberate blocks concur with the anticipated outcome of the button press responses. The longer response times for deliberate decision blocks also support the notion that participants categorically added more value to these decisions. The findings for the arbitrary blocks show a more random distribution in terms of button presses, as well as much shorter response times. This suggests that choices in these blocks may have been made haphazardly instead of deliberately. Participants were therefore effectively influenced by the distinction made between arbitrary and deliberate blocks. However, there was no statistically significant difference between the neurophysiological data of the respective decision types. A 2019 study conducted by Verbaarschot et al. found a similar result. They constructed an experiment where participants performed choice tasks in two separate computer game environments (an arbitrary environment and a deliberate environment). In these artificial environments, participants had to act to free a digital avatar. Upon analysis, they also found the presence of the intention to act arose before the awareness of the intention to do so across both arbitrary and deliberate decision classes (Verbaarschot et al., 2019).

It is important to note that a failed understanding of the decision types in the presented experiment would more likely result in both types of decisions being considered deliberate and not arbitrary. This is precisely because of the emotive component present in the choices. The content of the choices presented in this study comprises of the emotional and moral components of real-world choices. It can therefore be stated that the RP is present in the neurophysiological data of deliberate decision-making, where deliberate decisions are defined as choices with consequences.

As with many similar ERP studies, the moment of choosing is assumed to coincide with the button press event (Herrmann, et al., 2008; Maoz, et al., 2019). Even though this may introduce some latency in terms of the onset of the decision time, it is consistent across participants. More importantly, the experiment required participants to press the button the moment they made their choice, and since the build-up of the RP starts roughly 300 ms before the time-locked button press event, it can be assumed that the onset of the RP precedes the moment of conscious awareness of intent. The potential latency would only affect changes in the micro-scale temporal domain. Moreover, Maoz et al. showed that there was a clear discrepancy between the decision times of arbitrary and deliberate decisions (Maoz, et al., 2019). Similarly, participants in this study responded faster to arbitrary blocks than to deliberate blocks.

Other studies have previously suggested that the RP may be more indicative of the preparation to react (Alexander, et al., 2016) or the expectation to make a choice (Herrmann, et al., 2008) rather than the actual content of the choice, i.e. only indicative of a cortical build-up in anticipation of an executive decision task. However, the observed differences in ERPs between different participants, depending on their respective relationships to violent crime, demonstrated that the content of the decision may influence the RP and P300 amplitudes. Consequently, the RP may not be a neurophysiological phenomenon that simply arises because a choice task is present. The study suggests that personal experiences, specifically traumatic personal experiences, observably influence neurophysiological responses during the decision-making process. Although not statistically significant, these observations were evident for both the peak RP and peak P300 amplitudes. It was interesting to find that within the Crime I group the peak RP amplitude was observably increased while the peak P300 amplitude was observably decreased at electrode Fz for individuals with 1^st^ person exposure to violent crime. Electrode Fz records from the intentional and motivational centres of the brain (Teplan, 2002). The amplitude of the P300 peak may therefore resemble the emotional valence of a response, and a similar trend seen for the RP, may suggest that the RP too is influenced by response content. This finding, however, requires further research to confirm and will form part of future studies.

The study succeeded in showing that there were no neurophysiological differences between arbitrary and deliberate decisions, as the decision classes were defined at the onset of the study. The study also demonstrated that there were no neurophysiological differences between the convict and acquit trials or the left- and right-hand button presses. There was also a clear RP build-up prior to the button press events for both arbitrary and deliberate decisions. Furthermore, the choices presented in this research were more representative of choices with consequences than the choices presented in previous RP studies. Moreover, since the presented questions comprised of a moral component, the emotional context of participants’ responses could be evaluated. This served to establish a clear correlation between the peak amplitudes exhibited in scalp potentials and certain emotional triggers.

Although inconclusive with regards to the role conscious will has to play in deliberate choice, the study provides a glimpse into the delicate architecture underlying higher order, deliberate and emotional decision-making.

## 5. Limitations

It must be acknowledged that this study only considered a limited number of electrode sites. This study also focussed on specific ERPs. There might be value in expanding the scope of ERPs, as well as number of electrode sites, and implement machine learning algorithms to detect other neurophysiological regions and markers significant to deliberate decision-making. Moreover, since the Crime I: Y sample size was too small to have relevant statistical power, participants who fell into both the Crime I and Crime II groups could not be eliminated from consideration for either of the two group analyses. Therefore, to properly investigate the effect of personal exposure to violent crime on the neurophysiological markers of decision-making, a larger group of participants with 1^st^ person exposure to violent crime needs to be studied.

The data that support the findings of this study are available from the corresponding author upon reasonable request.

## 6. Acknowledgements

This research was funded by the Division of Biomedical Engineering at the University of Stellenbosch, South Africa.

## References

Alexander, P. et al., 2016. Readiness potentials driven by non-motoric processes. Consciousness and Cognition, Volume 39, pp. 38–47.

Batthyany, A., 2009. Mental Causation After Libet and Soon: Reclaiming Conscious Agency. In: A. Batthyany & A. C. Elitzur, eds. Irreducibly Conscious: Selected Papers on Consciousness. Heidelberg: Universitätsverslag Winter.

Bode, S. et al., 2014. Demystifying “free will”: The role of contextual information and evidence accumulation for predictive brain activity. Neuroscience & Biobehavioural Reviews, Volume 47, pp. 636–645.

Bonn, G., 2013. Re-conceptualizing free will for the 21st century: acting independently with a limited role for consciousness. Frontiers in Psychology, 920(4), pp. 1–9.

Bossearts, P. & Murawski, C., 2015. From behavioural economics to neuroeconomics to decision neuroscience: the ascent of biology in research on human decision making. Current Opinion in Behavioural Sciences, 2015(5), pp. 37–42.

Dawson, B. & Trapp, R. G., 2004. Basic & Clinical Biostatistics. Fourth ed. New York: MCGraw Hill.

Herrmann, C. S. et al., 2008. Analysis of a choice-reaction task yields a new interpretation of Libet’s experiments. International Journal of Psychology, 67(2), pp. 151–157.

Jo, H. G. et al., 2013. Spontaneous EEG fluctuations determine the readiness potential: is preconscious brain activation a preparation process to move?. Experimental Brain Research, 231(4), pp. 495–500.

Kornhuber, H. & Deecke, L., 1990. Readiness for movement - the Bereitschaftspotential Story. Current Contents Life Sciences, Volume 33, p. 14.

Kornhuber, H. H. & Deecke, L., 1965. Changes in the brain potential in voluntary movements and passive movements in man: Readiness pontential and reafferent potentials. Pflugers Archiv fur die Gesamte Physiologie des Menschen und der Tiere, Volume 284, pp. 1–17.

Lavazza, A., 2016. Free Will and Neuroscience: From Explaining Freedom Away to New Ways of Operationalizing and Measuring It. Frontiers in Human Neuroscience, Volume 10, p. 262.

Lei, X. & Liao, K., 2017. Understanding the Influences of EEG Reference: A Large-Scale Brain Network Perspective. Frontiers in Neuroscience, Volume 11, p. 205.

Libet, B., Gleason, C. A., Wright, E. W. & Pearl, D. K., 1983. Time of conscious intention to act in relation to onset of cerebral activity (readiness-potential) the unconscious initiation of a freely voluntary act. Brain, 106(3), pp. 623–642.

Maoz, U., Yaffe, G., Koch, C. & Mudrik, L., 2019. Neural precursors of decisions that matter - an ERP study of deliberate and arbitrary choice. eLife, 2019(8), p. e39787.

Noordzij, M. et al., 2010. Sample size calculations: basic principles and common pitfalls. Nephrology Dialysis Transplantation, 25(2010), pp. 1388–1393.

Peirce, J. W., 2009. Generating stimuli for neuroscience using PsychoPy. Frontiers in Neuroinformatics, 2(10).

SAS Institute Inc., 2011. Applied Analytics Using SAS Enterprise Miner Course Notes. Cary, NC, USA: SAS Institute Inc.

SAS Institute Inc., 2017. SAS/STAT 14.3 User’s Guide. Cary, NC, USA: SAS Institute Inc.

Soon, C., He, A., Bode, S. & Haynes, J., 2013. Predicting free choices for abstact intentions. Proceedings of the National Academy of Sciences of the United States of America, 110(15), pp. 6217–6222.

Soon, C. S., Brass, M., Heinze, H.-J. & Haynes, J.-D., 2008. Unconscious determinants of free decisions in the human brain. Nature Neuroscience, Volume 11, pp. 543–545.

Teplan, M., 2002. Fundamentals of EEG Measurement. Mesurement Science Review, 2(2).

Verbaarschot, C., Farquhar, J., Haselager, P., 2015. The search for intentions and Readiness Potentials. Consciousness & Cognition, Volume 33, p. 300–315.

Verbaarschot, C., Farquhar, J., Haselager, P., 2019. Free Wally: Where motor intentions meet reason and consequence. Neuropsychologia, Volume 133, p. 107156.

Wolpe, N. & Rowe, J. B., 2014. Beyond the “urge to move”: objective measures for the study of agency in the post-Libet era. Frontiers in Human Neuroscience, Volume 8, p. 450.

